# High-Resolution Genomic Profiling of Carbapenem-Resistant *Klebsiella pneumoniae* Isolates: A Multicentric Retrospective Indian Study

**DOI:** 10.1101/2021.06.21.449240

**Authors:** Geetha Nagaraj, Varun Shamanna, Vandana Govindan, Steffimole Rose, D. Sravani, K. P. Akshata, M.R. Shincy, V.T. Venkatesha, Monica Abrudan, Silvia Argimón, Mihir Kekre, Anthony Underwood, David M Aanensen, K. L. Ravikumar, The NIHR Global Health Research Unit for the Genomic Surveillance of Antimicrobial Resistance

## Abstract

**Background:** Carbapenem-resistant *Klebsiella pneumoniae* (CRKP) is a threat to public health in India due to its high dissemination, mortality, and limited treatment options. Its genomic variability is reflected in the diversity of sequence types, virulence factors, and antimicrobial resistance (AMR) mechanisms. This study aims to characterize the clonal relationships and genetic mechanisms of resistance and virulence in CRKP isolates in India.

**Materials and Methods:** We characterized 344 retrospective *K. pneumoniae* clinical isolates collected from 8 centers across India collected in 2013-2019. Susceptibility to antibiotics was tested with VITEK 2. Capsular types, MLST, virulence genes, AMR determinants, plasmid replicon types, and a single-nucleotide polymorphism (SNP) phylogeny were inferred from their whole genome sequences.

**Results:** Phylogenetic analysis of the 325 *Klebsiella* isolates that passed QC revealed 3 groups: *K. pneumoniae sensu stricto* (n=307), *K. quasipneumoniae* (n=17), and *K. varicolla* (n=1). Sequencing and capsular diversity analysis of the 307 *K. pneumoniae sensu stricto* isolates revealed 28 sequence types, 26 K-locus types, and 11 O-locus types, with ST231, KL51, and O1V2 being predominant. *bla*_OXA-48-like_ and *bla*_NDM-1/5_ were present in 73.2% and 24.4% of isolates respectively. The major plasmid replicon types associated with carbapenase genes were IncF (51.0%), and Col group (35.0%).

**Conclusion:** Our study documents for the first time the genetic diversity of K- and O-antigens circulating in India. The results demonstrate the practical applicability of genomic surveillance and its utility in tracking the population dynamics of CRKP. It alerts us to the urgency for longitudinal surveillance of these virulent and transmissible lineages.

**summary:** We report insights into genome sequences of Indian *K. pneumoniae* isolates, highlighting the presence of high-risk international clones and genetic pools different from those predominating in other regions. Identification of multidrug-resistant and hypervirulent *K. pneumoniae* elicits public health concerns.

## INTRODUCTION

*Klebsiella pneumoniae* is a common cause of nosocomial and community-acquired infections in newborns, the elderly, and immunocompromised patients [1]. The ability of *K. pneumoniae* to adapt by gene transfer, their virulence, and the convergence of resistance in them have led to the emergence of strains causing severe and untreatable invasive infections. Carbapenem-resistant *Klebsiella pneumoniae* (CRKP) is one of the most important and challenging pathogens causing infections with high mortality [2]. Co-occurrence of multidrug-resistance (MDR) and hypervirulence among *K. pneumoniae* lineages has elicited concerns from a public health standpoint [3].

Epidemic lineages of CRKP have increasingly emerged and spread through global health care systems since they were first identified in 2001 [4]. The worldwide spread of these strains is alarming, as they are multidrug-resistant, with resistance to β-lactam antibiotics, fluoroquinolones, and aminoglycosides [5]. The resistance of these species is generally due to the production of carbapenemases, such as the Class A serine carbapenemase *Klebsiella pneumoniae* (KPC) and metallo-β-lactamases. Other causes include combinations of outer-membrane permeability loss and extended spectrum β-lactamase production [6].

According to the Center for Disease Dynamics, Economics and Policy, India has seen an increase in carbapenem resistance in *K. pneumoniae*, from 24% in 2008 to 59% in 2017, though several single-center studies have shown variable rates [7, 8, 9]. This dramatic increase in recent years is attributed, among other factors, to the lack of adequate medical intervention, prolonged hospitalization, the presence of comorbidities, and the overuse of antibiotics. The rate of mortality among patients with CRKP bloodstream infections is as high as 68% [10]. Despite the high disease burden, there are limited reports from India on the resistance mechanisms in MDR *K. pneumoniae* isolates, highlighting the need for comprehensive epidemiological surveillance results [10].

The expansion and dissemination of CRKP at different geographical locations throughout India necessitates a targeted analysis of the population structure, genomic mechanisms of resistance, and virulence of strains collected from the area of interest. Recent developments in understanding population structure highlight enormous genomic diversity and provide a basis for the pathogen to be tracked [11]. Whole genome sequencing (WGS) is a powerful tool for the characterization and surveillance of pathogens. It offers an unparalleled opportunity to explore genomic content, and verify and evaluate the diversity of clusters and the frequency of contemporary isolates. It is now well-positioned to become the gold standard to resolve the knowledge gap [12].

Understanding of the mechanism by which this bacterium causes various infections is still basic, and most studies have limitations due to the narrow range of virulence factors being investigated. Little is known about the epidemiology of the capsule because serological and molecular typing are not commonly available, and several isolates are not typable using these methods. Phenotypic studies have identified 77 distinct capsule types (K-types) and genomic studies have identified 134, but the true extent of capsule diversity remains unknown [13]. WGS can provide new insights into disease transmission, virulence, and AMR dynamics when combined with epidemiological, clinical, and phenotypic microbiological information [12].

Since 2013, the Central Research Laboratory, Kempegowda Institute of Medical Sciences (CRL, KIMS) in India has developed a network of tertiary care hospitals, medical college hospitals, and stand-alone diagnostic laboratories across India. This network was extended for collection of retrospective isolates belonging to World Health Organization (WHO) priority bacterial pathogens. In this report, we characterize the clonal relationships and genetic mechanisms of resistance and virulence in CRKP isolates in India. WGS was performed on a retrospective collection of 344 *K. pneumoniae* isolates to characterize their relationships, multilocus sequence type (MLST), capsular type, virulence genes, and AMR determinants.

## MATERIALS AND METHODS

### Bacterial Isolates and Phenotypic Characterization

The study included 344 retrospective (2013-2019) invasive and non-invasive putative *K. pneumoniae* isolates received from 8 teaching hospitals in 6 Indian regions. They were characterized at CRL, KIMS using the VITEK 2 (Biomeurieux) compact system. The results were interpreted according to the 2019 CLSI guidelines. Isolates with phenotypic carbapenem-resistance of resistant (R) or intermediate (I) are considered resistant (R). An isolate is designated MDR when it shows resistance to >=1 agent in >= 3 antimicrobial categories [12].

### Sequencing and Genomic Analyses

Genomic DNA was extracted from bacterial isolates with Qiagen QIAamp DNA Mini kit, in accordance with the manufacturer’s instructions. Double-stranded DNA libraries with 450 bp insert size were prepared and sequenced on the Illumina platform with 150 bp paired-end chemistry. The genomes of 325 samples that passed sequence quality control were assembled using Spades v3.14 to generate contigs and annotated with Prokka v1.5 [15, 16]. SNPs were identified for 307 *K. pneumoniae* isolates by mapping of reads to the NCBI reference genome, *K. pneumoniae* strain NTUH-K2044, NC_006625.1 (Supplementary Table 1). MLST, K-and O-antigen types, virulence and AMR genes were identified using the protocols as detailed in www.protocols.io [17].

## RESULTS AND DISCUSSION

### Summary of the Collection

The collection consisted of 325 isolates from patients aged 7 days to 96 years, of whom 60% were 50-80 years old. Most isolates were from urine (31.4%) and blood (29.8%) (Supplementary Table 2). VITEK 2 Compact was used to identify 325 isolates as *K. pneumoniae*, and these were reassigned to the species *K. pneumoniae* (n=307, 94.4%), *K. quasipneumoniae* (n=17, 5.5%), and *K. variicola* (n=1, 0.1%) by sequencing. This highlights the limitation of traditional identification methods to distinguish species within the *K. pneumoniae* species complex. In addition, the use of genomic tools unmasks the true clinical significance of each phylogroup and their potential epidemiological specificities [15].

### Clonal Distribution

ST231 was the most common sequence type (n = 107, 34.8%), followed by ST147 (n = 73, 23.5%), and ST14 (n=26, 8.5%), accounting for 67.1% of total *K. pneumoniae* isolates (Table 1, Supplementary Figure 1). High prevalence of ST231 is in concordance with published data from India [16]. ST258 is recognized as the most prevalent and extensively disseminated KPC-producing *K. pneumoniae* in many countries, which made its absence in our collection noteworthy. ST11, a single-locus variant of ST258 and a prevalent clone associated with the spread of KPC in Asia (particularly in China and Taiwan), was identified in 1.3% of the *K. pneumoniae* isolates [18]. ST147 and ST14, described as international high-risk clones associated with extensive drug resistance (XDR), accounted for 23.5% and 8.5% of the isolates in this study, respectively [19, 20]. One novel ST (ST5603) with XDR was identified in a single *K. pneumoniae* isolate (Supplementary Table 2). The novel ST (gapA2-infB1-Mdh1-Pgi8-phoE10-rpoB4-tonb202) was a single-locus variant of ST890, varying at the tonb gene (tonb61), and possessed the KL107 and O1v1 loci.

**Table 1.**
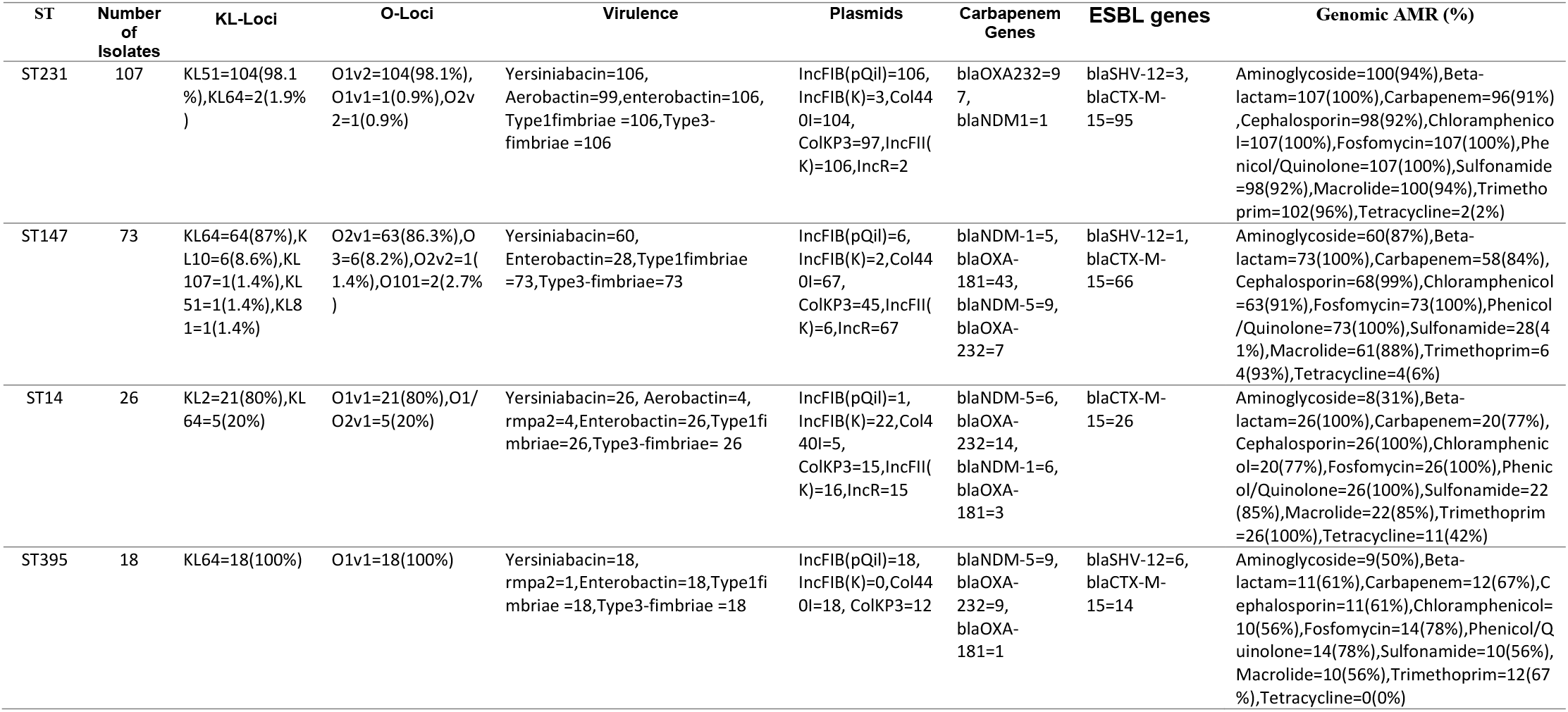
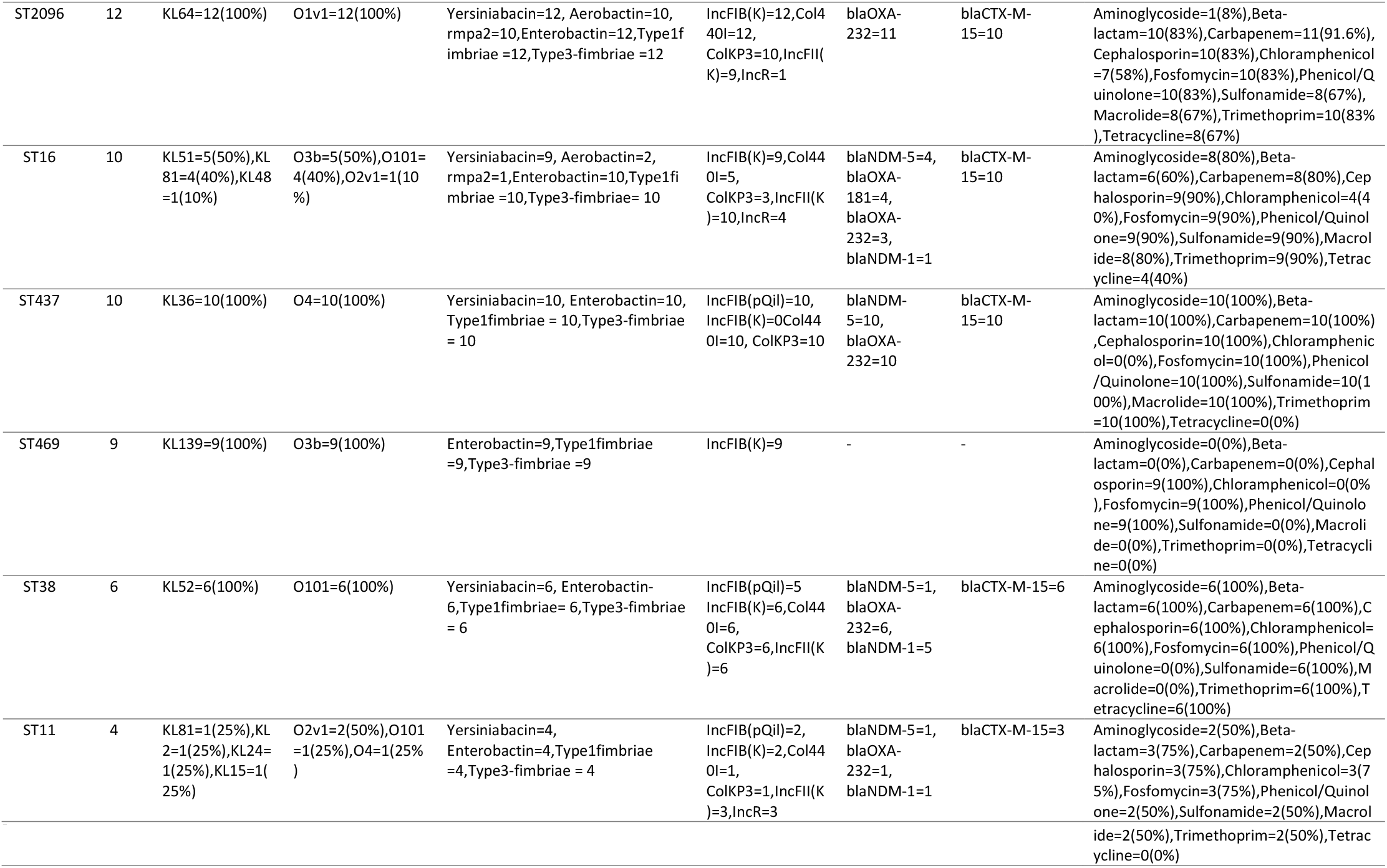
Distribution of AMR and Virulence genes across top five STs prevalent in *K.pneumoniae* isolates from India.

KL51 (n=111/307, 36.1%) and KL64 (n=104/307, 33.8%) were dominant K-loci types in this study. The phylogenetic tree of the 307 isolates shows the correlation between capsular locus type and sequence type, for example of KL51 with ST231, and of KL64 with ST147 and ST395 (Supplementary Figure 1). Notably, no isolates were assigned to KL1, despite its high prevalence in a previous study [21].

Out of 11 O-serotypes identified, O1, O2 and O3 together accounted for 90.2% of 307 *K. pneumoniae* samples (Supplementary Table 1). Strikingly, the O3b locus which is considered to be rare, was detected in 5.2% (16/307) of the isolates [21]. Other serotypes present included O4 (4.5%), OL101 (4.5%), OL103 (0.33%), and O5 (0.33%). O-locus (OL) types also showed ST-specific distribution. In particular, O1v1is found in STs 101, 1322, 14, 15, 2096, 231, 2497, 35, 48, and novel ST, while O2v2 is found in STs 307, 1248, 147, and 231. The stratification of O-types by patient age established that O1, O2, and O3 account for 81% of samples in the age group ≤5yrs (Supplementary Table 4).

### Virulence

More than 10 virulence factors account for the pathogenesis of CRKP, and their detection helps understand the pathogenesis of different strains [22]. In the present study, a total of 33 genes belonging to 6 major virulence factors were observed. The core virulence genes Type I and III fimbriae, enterobactin, AcrAB efflux pump, and regulators (RmpA, RcsAB) were identified in all 307 isolates. The acquired virulence genes coding for colibactin, nutrition factor, salmochelin, and rmpA were completely absent. Yersiniabactin, an iron uptake locus (ybtAEPQSTUX), was identified in 89.9% (276/307) of the isolates, and the regulatory gene rmpA2 was present in 5.5% (17/307). Aerobactin, a critical virulence factor for Hypervirulent *K. pneumoniae* (HvKP), was found in 38.1% (n=117) of the isolates, all belonging to ST231 or ST2096 (Supplementary Figure 3). Of the *K. pneumoniae* ST231-KL51 isolates, 94% were characterized by a virulence score of 4 (Supplementary Table 1). Phylogenetic analysis, using the dataset from this study and a global collection of ST231 isolates, identified a sublineage that has acquired aerobactin and yersiniabactin, as well as the OXA-232 carbapenemase [23].

### Resistance Profile and Their Distribution

Accumulation of AMR in *K. pneumoniae* is primarily due to horizontal gene transfer (HGT) aided by plasmids and mobile genetic elements. Since the first report of CRKP in 1996, the incidence of this MDR pathogen has increased significantly. The resistance is primarily due to production of acquired carbapenemases *bla*_KPC_, *bla*_OXA_, *bla*_NDM_, and the combinatorial mechanism of ESBL activity with loss of outer membrane porins [24]. This has become worrisome particularly at a time when no new promising antimicrobial agents are on the horizon [25]. For public health initiatives, understanding their emergence and distribution over a diverse geographical region is needed [26].

307 *K. pneumoniae* isolates displayed phenotypic resistance to Ampicillin (99%), Gentamicin (78.8%), Amikacin (73%), Amoxicillin-clavulanate (91.9%), Piperacillin-tazobactam (93.5%), Cefepime (92.2%), Ceftriaxone (91.9%), Ciprofloxacin (95.4%), Carbapenems (91.2%), Cotrimoxazole (87.9%), and Colistin (8.8%). We observed 85 different resistance profiles in *K. pneumoniae* isolates (Supplementary Table 3). Of the 307 isolates, 280 were carbapenem-resistant. The majority of these showed phenotypic resistance to both meropenem and imipenem (256/280, 91.4%, Figure 1A), explained by the presence of *bla*_OXA-48-like_ genes, *bla*_NDM_ genes, and a combination of ESBLs (*bla*_CTX-M-15_, *bla*_CTX-M-71_, *bla*_SHV-12_, *bla*_SHV-13_, *bla*_SHV-27_, *bla*_SHV-31_, *bla*_SHV-41_) and inactivation of porins Ompk35/36 [27]. Of the carbapenem-resistant isolates, 80.3% (225/280) carried *bla*_OXA-48-like_ genes (*bla*_OXA181_ and *bla*_OXA232_), and 26.7% (75/280) carried *bla*_NDM-1/5_, which supports the reported changing trends in carbapenem resistance [25]. Of the 225 isolates with *bla*_OXA48-like_ genes, 60.7% (170/280) carried *bla*_OXA232_, and 19.6% (55/280) carried *bla*_OXA181_. *bla*_OXA232_ was present mostly within ST231 (97/170) in line with a previous report of widely dissemination of carbapenem-resistant ST231 with *bla*_OXA232_ across India (Figure 1B) [16]. Notably, no *bla*_KPC_ genes were detected, which are the predominant carbapenemases found in Europe and in Colombia [28, 29].

**Figure 1.**
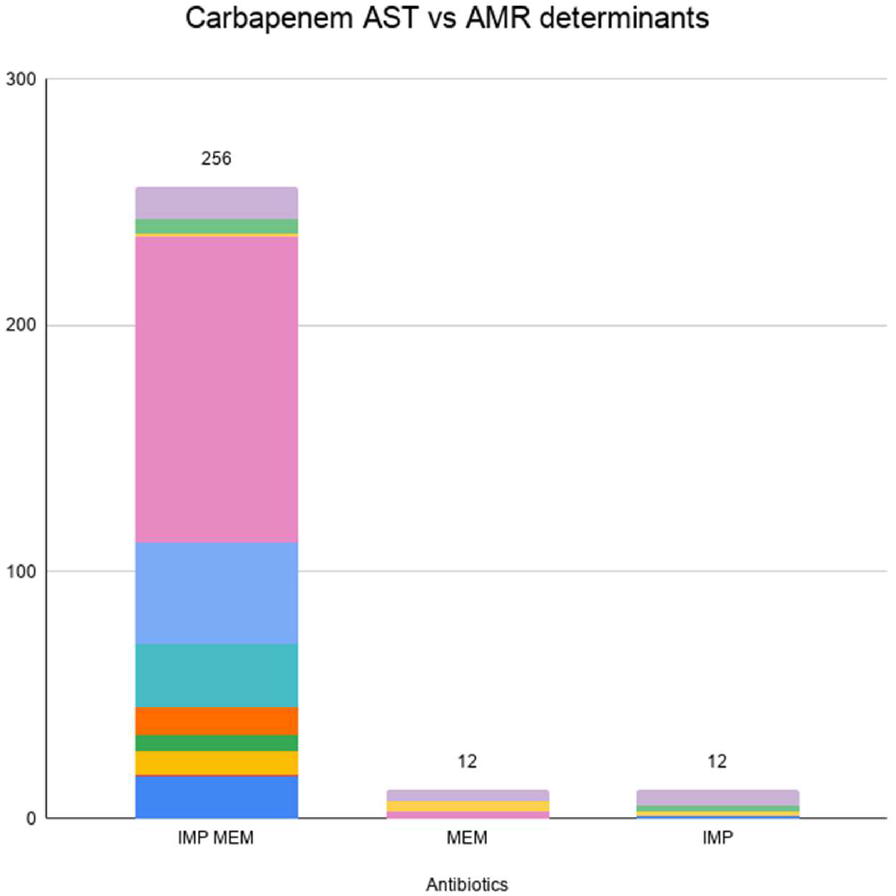
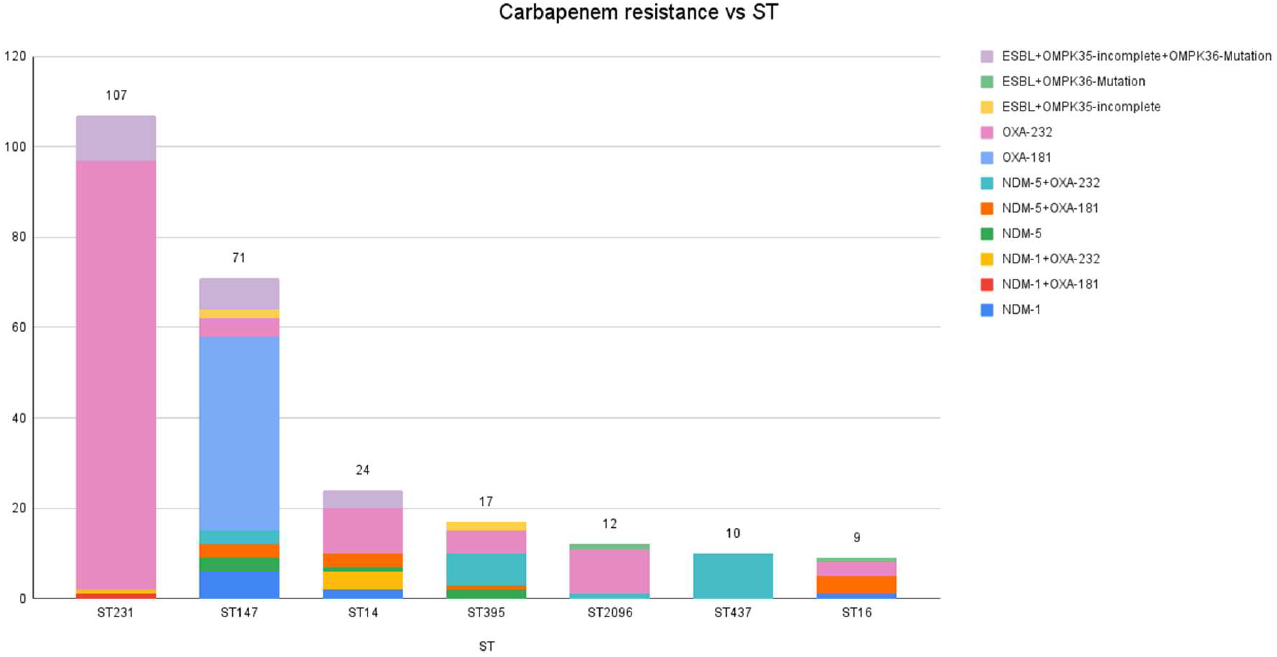
Carbapenem resistance mechanisms by ST. A) Mechanisms of resistance to carbapenems identified in the genomes of 280 isolates grouped by their phenotypic carbapenem resistance profile. Genes responsible for carbapenems are indicated. B) The determinants present in each ST and responsible for carbapenem resistance. Note, where more than one determinant is present, the single determinant or combination which is likely to result in the highest increased in susceptibility to carbapenems is recorded.

In the present dataset, resistance to carbapenems is also conferred by ESBLs when combined with porin loss-of-function mutations [30]. Disruption of the Ompk35 porin (n=231) was observed in the study isolates, and insertion mutations of TD/GD amino acids at position 115 of Ompk36 gene were found (n=224). The disruption of major outer membrane protein genes (Ompk35/Ompk36) along with ESBLs (*bla*_CTX-M-15_, *bla*_CTX-M-71_, *bla*_SHV-12_, *bla*_SHV-13_, *bla*_SHV-27_, *bla*_SHV-31_, *bla*_SHV-41_) were detected in 92.1% of the isolates (258/280). The presence of multiple ESBL genes in a single isolate (n=12), highlights the emerging complexity of antibacterial resistance repertoire.

### Plasmid Repertoire

The predominant plasmid replicons present in the 307 *K. pneumoniae* isolates are Col440I (79.2%), ColKP3 (68.7%), and IncFII (K) (57.0%) (Supplementary Figure 4). Other plasmid replicons detected are IncX, IncH, IncL, IncA/C, IncY, and IncQ. Understanding such vectors carrying the AMR genes could help in improving strategies to control AMR dissemination.

In the collection, 225 isolates carried *bla*_OXA48-like_ genes. We observed that 93.7% (n=211) of them harbored a plasmid with the ColKP3 replicon sequence (Supplementary Figure 2). A similar association between the ColKP3 replicon and the *bla*_OXA-232_ gene was also observed in ST231 isolates from around the world (Supplementary Figure 5) [23].

### High-risk clone – ST231

ST231, an emerging CRKP epidemic clone, was reported from India, France, Singapore, Brunei, Darussalam, and Switzerland [31]. In south-east Asia, this clone was found to be MDR, combining resistance to carbapenemes, extended-spectrum cephalosporins, and broad-spectrum aminoglycosides [32]. In India, ST231 was first reported in 2013 in Delhi, with the isolate carrying *bla*_OXA-232_ as the predominant *bla*_OXA48-like_ carbapenemase variant [16].

Isolates belonging to ST231 (n=107) were entirely genotypically MDR. Of the ST231 isolates, 91.5% (n=98/107) carried *bla*_OXA-232_ genes, and 99.0% (n=106/107) carried porin (Ompk35/Ompk36) mutations and ESBL genes (*bla*_CTX-M-15_, *bla*_SHV-12_), a possible mechanism for non-susceptibility to carbapenems (Figure 2). In addition, ST231 genomes harbored other resistance genes, including rmtF (39/107, aminoglycoside resistance), ermB (98/107, macrolide resistance), catA1 (104/107, phenicol resistance), sul1 (101/107, sulfonamide resistance), dfrA12 (101/107, trimethoprim resistance), as well as mutations in gyrA_83I and parC_80I (107/107, fluoroquinolone resistance).

**Figure 2.**
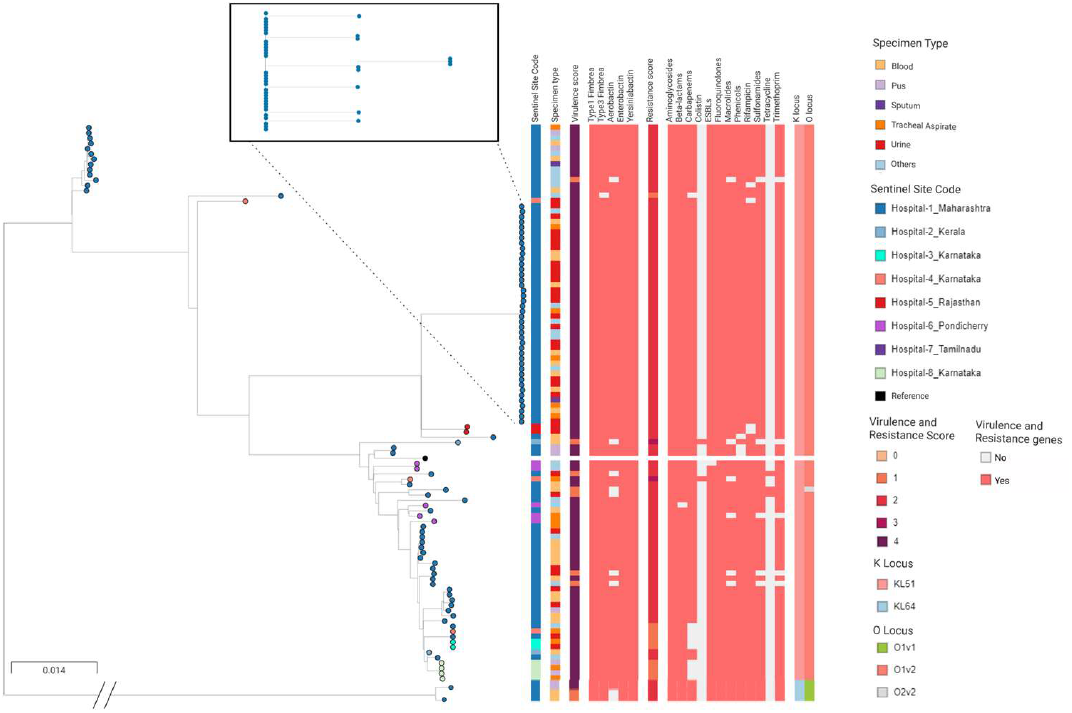
Phylogenetic analysis of ST231 lineage. A phylogenetic tree of 107 ST231 genomes from India. The lineage highlighted in the box is shown in detail, with metadata blocks showing ST, K-locus, AMR determinants and virulence factors. The full data are available at https://microreact.org/project/5GDXCkZsvajcXTmW89LRED/746bb511.

When compared to global genomes, the ST231 genomes from our study and those from a previous study in India (BioProject accession number PRJEB30913) shared a common repertoire of AMR and virulence determinants, and were found interspersed with (and often basal to) genomes from other countries in the tree, thus pointing to international dissemination (Supplementary Figure 5) [23, 33]. The tree of 107 ST231 isolates from this study showed evidence of clonal spread of ST231 carrying *bla*_OXA-232_ and *bla*_CTX-M-15_ within one hospital over a period of 3 years (Hospital 1, Figure 2). Forty-two (42) isolates formed a tight cluster with a mean SNP difference of 1 (range 0-3), and they were separated by at least 43 SNP differences from the remaining 65 ST231 isolates in this study. The isolates in this cluster were mostly from inpatients (37/42) and characterized by a median patient age of 73.5 years (range 26-89) compared to 67 years (7d-96) for all 239 *K. pneumoniae* isolates collected by this hospital. Altogether, this reveals a persistent outbreak of carbapenem and cephalosporin-resistant ST231 within Hospital 1 and underscores the need to strengthen infection prevention and control. Importantly, the tree also shows other ST231 genomes from Hospital 1 that show similar resistance and virulence profiles, but that can be clearly distinguished from this large outbreak by their clustering, highlighting the utility of WGS to rule cases out of an outbreak investigation even when other phenotypic or genotypic markers would be inconclusive.

We conclude that the MDR ST231 lineage carrying both important resistance and virulence determinants is a major and rapidly disseminating CRKP high-risk clone in India capable of causing nosocomial outbreaks [16]. The emergence of the ST231 clonal lineage has also been reported in Switzerland, France, and Thailand recently [34–36]. The presence of MDR and virulence genes poses a risk in that the lineage may be a reservoir of virulence-associated genes that can be passed on by HGT to other lineages. This means that a high level of vigilance and monitoring is required. As shown by other studies, WGS can be used as an outbreak detection tool, allowing the detection of widely dispersed outbreaks that might not be otherwise identified.

## CONCLUSION

The study establishes the presence of several high-risk MDR CRKP clones in clinical samples collected across India. It represents the basis for genomic surveillance of emerging CRKP in India, providing critical information that can be used to track the emergence and dissemination, and assess the potential impact, of important variants. The lack of structured surveillance framework and inability to access patient clinical data has limited our interpretation. To the best of our knowledge, this is the first WGS study from India to document genetic diversity of K-and O-antigens circulating in Indian CRKP isolates.

## Abbreviations

KPC: Class A serine carbapenemase *Klebsiella pneumoniae*
MDR: multidrug-resistant
CRKP: Carbapenem-resistant *Klebsiella pneumoniae*
WGS: Whole genome sequencing
MLST: multilocus sequence type
ST: sequence type
HvKP: Hypervirulent *K. pneumoniae*
HGT: horizontal gene transfer

## ACKNOWLEDGMENTS

Members of the NIHR Global Health Research Unit for the Genomic Surveillance of Antimicrobial Resistance: Khalil Abudahab, Harry Harste, Dawn Muddyman, Ben Taylor, Nicole Wheeler, and Sophia David of the Centre for Genomic Pathogen Surveillance, Big Data Institute, University of Oxford, Old Road Campus, Oxford, United Kingdom and Wellcome Genome Campus, Hinxton, UK; Pilar Donado-Godoy, Johan Fabian Bernal, Alejandra Arevalo, Maria Fernanda Valencia, and Erik C. D. Osma Castro of the Colombian Integrated Program for Antimicrobial Resistance Surveillance – Coipars, CI Tibaitatá, Corporatión Colombiana de Investigación Agropecuaria (AGROSAVIA), Tibaitatá – Mosquera, Cundinamarca, Colombia; K.N Ravishankar of the Central Research Laboratory, Kempegowda Institute of Medical Sciences, Bengaluru, India; Iruka N Okeke, Anderson O. Oaikhena, Ayorinde O. Afolayan, Jolaade J Ajiboye, and Erkison Ewomazino Odih of the Department of Pharmaceutical Microbiology, Faculty of Pharmacy, University of Ibadan, Oyo State, Nigeria; Celia Carlos, Marietta L. Lagrada, Polle Krystle V. Macaranas, Agnettah M. Olorosa, June M. Gayeta, and Elmer M. Herrera of the Antimicrobial Resistance Surveillance Reference Laboratory, Research Institute for Tropical Medicine, Muntinlupa, the Philippines; Ali Molloy, alimolloy.com; John Stelling, The Brigham and Women’s Hospital; and Carolin Vegvari, Imperial College London.

