## Supplementary table for "High-Resolution Genomic Profiling of Carbapenem-Resistant *Klebsiella pneumoniae* Isolates: A Multicentric Retrospective Indian Study"

[a] Members of the NIHR Global Health Research Unit on Genomic Surveillance of Antimicrobial Resistance are listed in the Acknowledgments.

***Index***

Supplementary Table 1: In Excel file

Supplementary Table 2: This document, page 2.

Supplementary Table 3: This document, page 3.

Supplementary Table 4: This document, page 8.

Supplementary Figure 1: This document, page 9.

Supplementary Figure 2: This document, page 10.

Supplementary Figure 3: This document, page 11.

Supplementary Figure 4: This document, page 12.

Supplementary Figure 5: This document, page 13.

### Supplementary Tables

**Supplementary Table 2.** The *K.pneumoniae* collection in this study by collection year, age, gender, clinical manifestation, phenotypic AMR and Genotypic AMR.

(In Excel file.)

**Supplementary Table 2.** Demographic and clinical characteristics of 325 *K. pneumoniae* isolates.

| **Characteristic** | **No. Isolates** |
|---|---|
| **GENDER** | |
| MALE | 179 |
| FEMALE | 146 |
| **AGE (in years)** | |
| <=2 | 11 |
| >2 - <=10 | 2 |
| >10 - <=20 | 13 |
| >20- <=30 | 14 |
| >30 - <=40 | 19 |
| >40 - <=50 | 25 |
| >50 - <=60 | 50 |
| >60 - <=70 | 79 |
| >70 - <=80 | 66 |
| >80 - <=90 | 41 |
| >90 - <=100 | 5 |
| | 325 |
| **Specimen origin** | |
| Community-acquired | 0 |
| Hospital-acquired | 325 |
| **Submitted As** | |
| Resistant to at least to one class of antibiotic | 325 |
| Carbapenem non-susceptible | 280 |
| **Specimen Type** | |
| Urine | 102 |
| Blood | 97 |
| Tracheal aspirate | 33 |
| Pus | 27 |
| Sputum | 13 |

| | |
|---|---|
| Wound | 11 |
| Tissue | 8 |
| Cathetar | 7 |
| Abdominal fluid | 3 |
| Abscess, abdominal | 2 |
| Abdomen | 2 |
| Aspirate | 2 |
| Broncho-alveolar lavage | 2 |
| Biopsy | 2 |
| Abcesses | 1 |
| Bile | 1 |
| Bone marrow | 1 |
| Bone | 1 |
| Catheter, central | 1 |
| Drain | 1 |
| Fluid | 1 |
| Liver | 1 |
| Pelvis | 1 |
| Mouth | 1 |
| Pacemaker | 1 |
| Rectal | 1 |
| Throat | 1 |
| | 325 |

**Supplementary Table 3.** Total number of *K. pneumoniae* isolates analyzed by the CRL during 2013 and 2019, isolates submitted for whole-genome sequencing, and high-quality *K. pneumoniae* genomes obtained, discriminated by sentinel site and AMR profile.

| | **Number of isolates** | | | | | | | |
|---|---|---|---|---|---|---|---|---|
| | **2013** | **2014** | **2015** | **2016** | **2017** | **2018** | **2019** | **Total** |
| **Total nos submitted for WGS** | 1 | 40 | 29 | 75 | 159 | 23 | 17 | 344 |
| High quality *K.pneumoniae* genomes | 1 | 35 | 29 | 65 | 141 | 22 | 14 | 307 |
| *By Sentinal sites* | | | | | | | | 0 |
| BAN | | | | | 14 | | 1 | 15 |
| JOD | | | | | 15 | 1 | | 16 |
| KER | | | 1 | 1 | 3 | | | 5 |
| KOL | | | 1 | | 4 | | | 5 |
| MAN | | | | 14 | 1 | | | 15 |
| MUM | 1 | 35 | 27 | 50 | 92 | 21 | 13 | 239 |

| | | | | | | | | |
|---|---|---|---|---|---|---|---|---|
| PON | | | | | 8 | | | 8 |
| TRI | | | | | 4 | | | 4 |
| *By AMR profile* | | | | | | | | |
| **AST profile** | **2013** | **2014** | **2015** | **2016** | **2017** | **2018** | **2019** | **Total Profile Count** |
| AMC AMK AMP CAZ CIP COL CSL CXA CXM FEP GEN IPM MEM NAL NIT SXT TZP | | | | | 2 | | | 2 |
| AMC AMK AMP CAZ CIP COL CSL CXA CXM FEP GEN IPM MEM NAL NIT TZP | | | | | 1 | | | 1 |
| AMC AMK AMP CAZ CIP CSL CXA CXM FEP GEN IPM MEM NAL NIT SXT TZP | | | | | 3 | | | 3 |
| AMC AMK AMP CAZ CIP CSL CXA CXM FEP GEN IPM MEM NAL SXT TZP NIT | | | | | 1 | | | 1 |
| AMC AMK AMP CFM CIP COL CRO FEP GEN IPM MEM SXT TZP | | | | 1 | 5 | 2 | | 8 |
| AMC AMK AMP CFM CIP COL CRO FEP GEN MEM SXT TZP IPM | | | 1 | 1 | 2 | | | 4 |
| AMC AMK AMP CFM CIP COL CRO FEP IPM MEM TZP | | 1 | | | | | | 1 |
| AMC AMK AMP CFM CIP CRO FEP GEN IPM MEM SXT TZP | | 19 | 20 | 21 | 49 | 8 | 7 | 124 |
| AMC AMK AMP CFM CIP CRO FEP GEN IPM MEM SXT TZP COL | | | 1 | | 1 | | | 2 |
| AMC AMK AMP CFM CIP CRO FEP GEN IPM MEM TZP | | | 1 | | 1 | | | 2 |
| AMC AMK AMP CFM CIP CRO FEP GEN MEM SXT TZP | | | | 1 | | | | 1 |
| AMC AMK AMP CFM CIP CRO FEP GEN MEM SXT TZP IPM | | 6 | | 7 | 7 | 1 | 2 | 23 |
| AMC AMK AMP CFM CIP CRO FEP GEN SXT TZP | | | | 1 | | | | 1 |
| AMC AMK AMP CFM CIP CRO FEP GEN SXT TZP MEM | | | | | 1 | | | 1 |
| AMC AMK AMP CFM CIP CRO FEP IPM MEM SXT TZP | | | | 1 | 3 | | 2 | 6 |
| AMC AMK AMP CFM CIP CRO FEP IPM MEM SXT TZP GEN | | | 1 | | 1 | | | 2 |
| AMC AMK AMP CFM CIP CRO FEP IPM MEM TZP | | 1 | | | | | | 1 |

| | | | | | | | | |
|---|---|---|---|---|---|---|---|---|
| AMC AMK AMP CFM CIP CRO FEP MEM SXT TZP IPM | | | 1 | | 1 | | 1 | 3 |
| AMC AMK AMP CFM CIP CRO FEP SXT TZP | | | | 2 | 2 | | | 4 |
| AMC AMK AMP CFM CIP CRO GEN MEM SXT TZP FEP | | | | | 1 | | | 1 |
| AMC AMK AMP CFM CIP CRO GEN SXT TZP FEP | | | | 1 | | | | 1 |
| AMC AMK AMP CIP COL CRO CSL CTX CXA ETP FEP GEN IPM MEM NAL NIT SXT TGC TZP | | | | 1 | | | | 1 |
| AMC AMK AMP CIP COL CRO CSL CXA CXM FEP GEN IPM MEM NAL NIT SXT TZP | | | | | 2 | | | 2 |
| AMC AMK AMP CIP COL CRO CSL CXA CXM FEP GEN IPM MEM NAL NIT TZP | | | 1 | | | | | 1 |
| AMC AMK AMP CIP COL CRO CSL CXA CXM FEP IPM MEM NAL NIT SXT TZP | | | | | 1 | | | 1 |
| AMC AMK AMP CIP CRO CSL CTX CXA ETP FEP GEN IPM MEM NAL NIT SXT TGC TZP | | | | 3 | | | | 3 |
| AMC AMK AMP CIP CRO CSL CTX CXA ETP FEP GEN IPM NAL NIT SXT TZP | | | | 1 | | | | 1 |
| AMC AMK AMP CIP CRO CSL CTX CXA ETP FEP IPM MEM NAL NIT SXT TGC TZP | | | | 2 | | | | 2 |
| AMC AMK AMP CIP CRO CSL CTX CXA ETP FEP IPM MEM NAL NIT SXT TGC TZP GEN | | | | 1 | | | | 1 |
| AMC AMK AMP CIP CRO CSL CTX CXA ETP FEP IPM NAL NIT SXT TGC TZP | | | | 1 | | | | 1 |
| AMC AMK AMP CIP CRO CSL CXA CXM ETP FEP GEN IPM MEM NAL SXT TZP | | | | | 1 | | | 1 |
| AMC AMK AMP CIP CRO CSL CXA CXM ETP FEP GEN IPM NAL NIT SXT TGC TZP | | | | | 3 | | | 3 |
| AMC AMK AMP CIP CRO CSL CXA CXM ETP FEP GEN NAL NIT SXT TGC TZP IPM | | | | | 1 | | | 1 |
| AMC AMK AMP CIP CRO CSL CXA CXM FEP GEN IPM MEM NAL NIT SXT TZP | | | | | 8 | | | 8 |

| | | | | | | | | |
|---|---|---|---|---|---|---|---|---|
| AMC AMK AMP CIP CRO CSL CXA CXM FEP GEN IPM MEM NAL SXT TZP NIT | | | | | 1 | | | 1 |
| AMC AMK AMP CIP CRO CSL CXA CXM FEP GEN IPM NAL NIT SXT TZP | | | | | 2 | | | 2 |
| AMC AMK AMP CIP CRO CSL CXA CXM FEP GEN IPM NAL NIT SXT TZP MEM | | | | | 2 | | | 2 |
| AMC AMP CAZ CIP CSL CXA CXM FEP IPM MEM NAL NIT TZP AMK | | | | | 1 | | | 1 |
| AMC AMP CAZ CSL CXA CXM FEP GEN IPM NAL NIT TZP | | | | | 1 | | | 1 |
| AMC AMP CFM CIP COL CRO FEP GEN IPM MEM SXT TZP | | | | 1 | 1 | | | 2 |
| AMC AMP CFM CIP COL CRO FEP GEN IPM MEM SXT TZP AMK | | | | | | 1 | | 1 |
| AMC AMP CFM CIP COL CRO FEP GEN MEM SXT TZP IPM | | | | 1 | | | | 1 |
| AMC AMP CFM CIP CRO FEP GEN IPM MEM SXT TZP | 1 | 2 | | 2 | 2 | | | 7 |
| AMC AMP CFM CIP CRO FEP GEN IPM MEM SXT TZP AMK | | 3 | | 2 | 1 | 1 | | 7 |
| AMC AMP CFM CIP CRO FEP GEN IPM MEM TZP | | 1 | | | 1 | | | 2 |
| AMC AMP CFM CIP CRO FEP GEN MEM SXT TZP AMK IPM | | | 1 | | 2 | | | 3 |
| AMC AMP CFM CIP CRO FEP GEN MEM SXT TZP IPM | | 1 | | 2 | 5 | | 1 | 9 |
| AMC AMP CFM CIP CRO FEP GEN SXT TZP IPM | | | | | 1 | | | 1 |
| AMC AMP CFM CIP CRO FEP IPM MEM SXT TZP | | | | | | 2 | | 2 |
| AMC AMP CFM CIP CRO FEP IPM MEM SXT TZP AMK | | | | | 1 | 1 | | 2 |
| AMC AMP CFM CIP CRO FEP IPM MEM SXT TZP GEN | | | | | 2 | | | 2 |
| AMC AMP CFM CIP CRO FEP IPM MEM TZP | | | | 1 | 1 | | | 2 |
| AMC AMP CFM CIP CRO FEP IPM MEM TZP GEN | | | | | 1 | 1 | | 2 |
| AMC AMP CFM CIP CRO FEP IPM MEM TZP GEN SXT | | | | | 1 | | | 1 |
| AMC AMP CFM CIP CRO FEP MEM SXT TZP AMK IPM | | | | | 3 | | | 3 |

| | | | | | | | | |
|---|---|---|---|---|---|---|---|---|
| AMC AMP CFM CIP CRO FEP MEM TZP IPM | | | | 2 | | 2 | | 4 |
| AMC AMP CFM CIP CRO FEP SXT TZP MEM | | | | | | 1 | | 1 |
| AMC AMP CFM CIP MEM TZP IPM | | | | 1 | | | | 1 |
| AMC AMP CIP COL CRO CSL CXA CXM IPM MEM NAL NIT TZP | | | | 1 | | | | 1 |
| AMC AMP CIP COL CSL CXA CXM IPM MEM NAL NIT TZP CRO | | | | | 1 | | | 1 |
| AMC AMP CIP CRO CSL CTX CXA ETP FEP IPM MEM NAL NIT SXT TGC TZP | | | | 1 | | | | 1 |
| AMC AMP CXA CXM SXT IPM NIT | | | | | 1 | | | 1 |
| AMC AMP FEP IPM MEM SXT TZP | | | | 1 | | | | 1 |
| AMK AMP CIP CRO CXA CXM FEP GEN NAL NIT PIP SXT AMC | | | | | 1 | | | 1 |
| AMK FEP GEN IPM MEM TZP AMP | | | | 1 | | | | 1 |
| AMP | | | | | 2 | | 1 | 3 |
| AMP CAZ CXA CXM NAL AMC | | | | | 1 | | | 1 |
| AMP CFM CIP CRO FEP GEN SXT TZP AMC AMK | | | | | | 1 | | 1 |
| AMP CFM CIP CRO FEP MEM SXT TZP AMC | | | 1 | | | | | 1 |
| AMP CFM CIP CRO IPM MEM AMC TZP | | 1 | | | | | | 1 |
| AMP CFM CIP CRO SXT AMC FEP TZP | | | | | 1 | | | 1 |
| AMP CIP CRO CSL CXA CXM FEP GEN NAL TZP AMC NIT | | | | | 1 | | | 1 |
| AMP CIP CRO CTX CXA GEN SXT AMC NIT TZP | | | | 1 | | | | 1 |
| AMP CIP CRO CTX CXA NIT SXT TGC | | | | 1 | | | | 1 |
| AMP CIP CRO CXA CXM FEP NAL SXT TZP AMC | | | | | 1 | | | 1 |
| AMP CIP CRO CXA CXM GEN NAL AMC | | | | | 1 | | | 1 |
| AMP CIP CRO CXA CXM GEN NAL SXT AMC NIT TZP | | | | | 1 | | | 1 |
| AMP CIP NAL NIT | | | | | 1 | | | 1 |
| AMP CIP NAL SXT AMC | | | | | 1 | | | 1 |

| | | | | | | | | |
|---|---|---|---|---|---|---|---|---|
| AMP CRO CXA CXM AMC | | | | | | 1 | | 1 |
| AMP CRO CXA CXM NAL SXT NIT | | | | | 1 | | | 1 |
| AMP NIT | | | | 2 | 1 | | | 3 |
| CAZ CIP DOR FEP GEN IPM LVX MEM TCC TZP CSL TGC | | | 1 | | | | | 1 |

**Supplementary Table 4.** Distribution of O-types in age group ≤ 5yrs.

| Patient_age | AgeGroup | Patient gender | K_locus | O_locus |
|---|---|---|---|---|
| 7d | 0-10 | Male | KL64 | O2v1 |
| 8d | 0-10 | Female | KL24 | O1v1 |
| 8d | 0-10 | Female | KL36 | O4 |
| 1m | 0-10 | Male | KL127 | OL101 |
| 1m | 0-10 | Female | KL2 | O1v1 |
| 3m | 0-10 | Male | KL21 | O3b |
| 1yr | 0-10 | Female | KL125 | O5 |
| 1yr | 0-10 | Male | KL2 | O2v1 |
| 1yr | 0-10 | Male | KL2 | O1v1 |
| 1yr | 0-10 | Male | KL51 | O1v2 |
| 1yr | 0-10 | Male | KL51 | O1v2 |
| 5yr | 0-10 | Male | KL52 | OL101 |

### Supplementary Figures

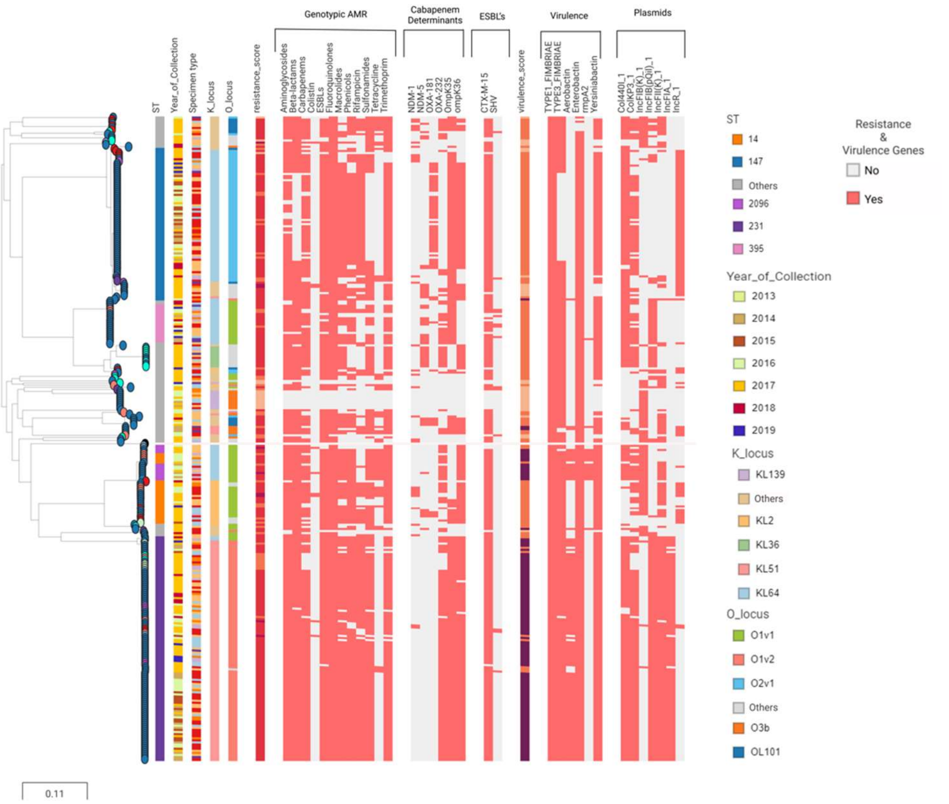

**Supplementary Figure 1.** Genomic surveillance of K.pneumoniae from India 2013-2019. Phylogenetic tree of 307 isolates from India inferred by iQTree. The tree leaves are coloured by sentinel site. Metadata blocks: Sequence Type, Year of collection, Specimen type, patient age & gender, Resistance score, resistance genes, Virulence score and genes, K-and O-loci. The data, including the full distribution of resistance determinants, are available at: https://microreact.org/project/w2WjcPV95L13PtYioDHAep/2ccbf0f3.

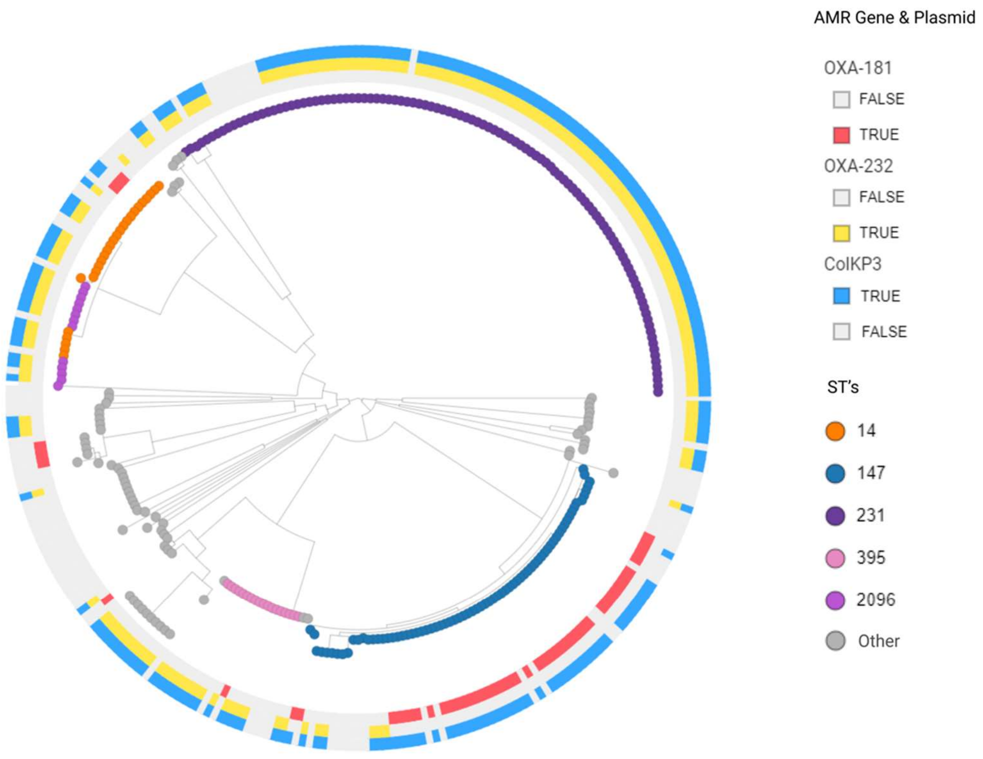

**Supplementary Figure 2.** MR figure showing association between ColKP3, OXA-232 and OXA-181.The full data are available at:

https://microreact.org/project/cpAqgWfKANmcoQarP3VCUk/f0b97011.

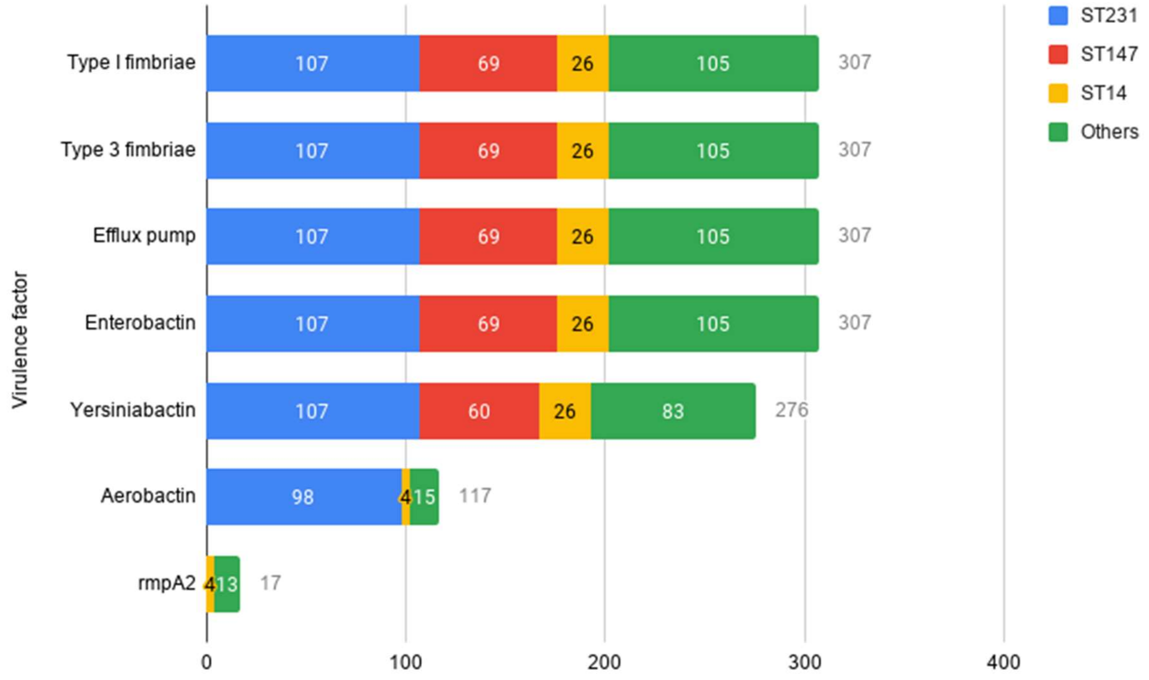

**Supplementary Figure 3.** Distribution of virulence genes in the collection of 307 genomes.

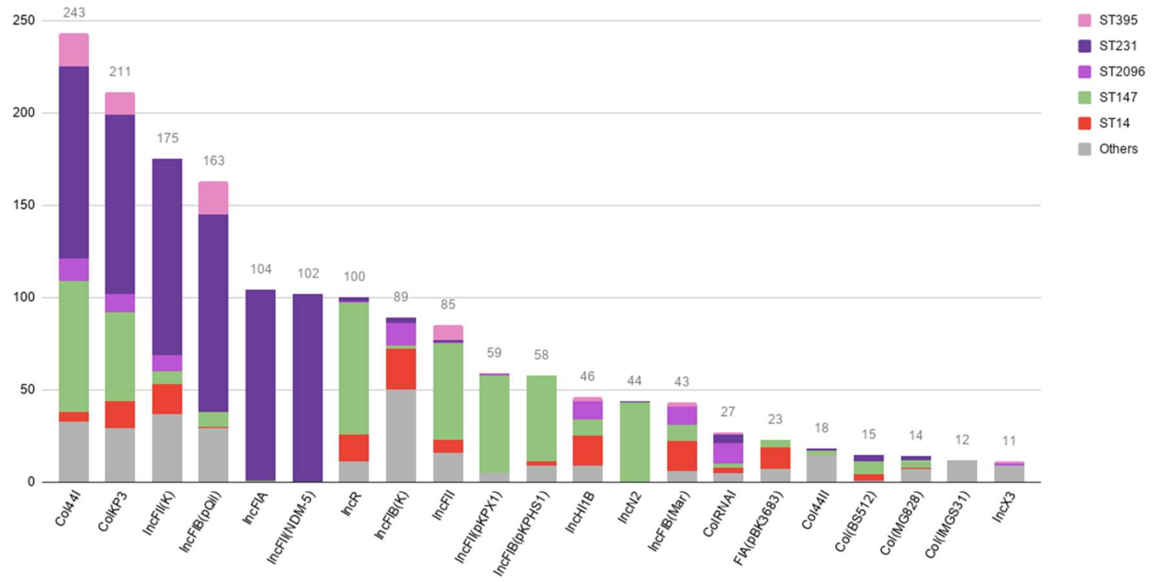

**Supplementary Figure 4.** Distribution of plasmids and their predominant STs in the collection of 307 genomes.

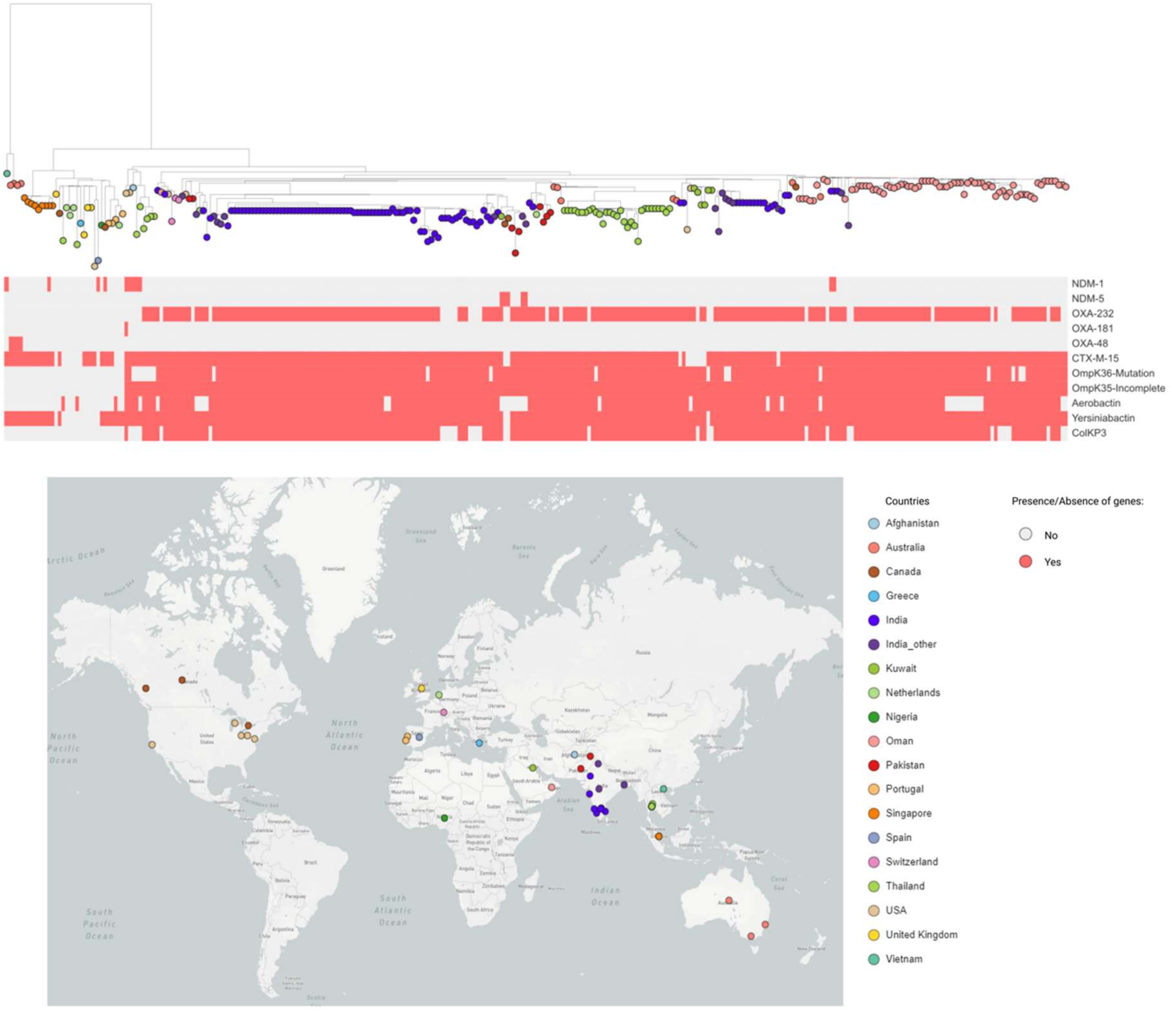

**Supplementary Figure 5.** Phylogenetic tree of 107 ST231 genomes was built against the global collection of ST231 (198) in pathogen watch. The tree leaves are coloured by countries as indicated on the map from Figure 3B. The tree blocks from Figure 3A represent the distribution of the carbapenemase genes and acquired resistance genes and mutations. The full data are available at: https://microreact.org/project/BNU6PXMPwWo4e8HtiZYtJ/dce19a20.
